## Supplemental Figures 1-7, supplemental Tables 1-4, Supplemental note 1 for "Characterisation of tumour microenvironment remodelling following oncogene inhibition in preclinical studies with imaging mass cytometry"

**Supplementary Table 1. Fluorophore-conjugated antibodies used for optimisation and validation of the antibody panel.**

| target[1] | clone | fluorophore | source | catalogue number |
|---|---|---|---|---|
| CD45 | 30-F11 | APC | Biolegend | 103112 |
| alphaSMA | 1A4 | Cy3 | Sigma-Aldrich | C6198 |
| MHC-II | M5/114.15.2 | APC | Biolegend | 107613 |
| vimentin | D21H3 | | | |
| PECAM/CD31 | 390 | APC | Biolegend | 102409 |
| F4/80 | CI:A3-1 | RPE | BioRad | MCA497PET |
| CD68 | FA-11 | PE/Cy7 | BioLegend | 137015 |
| EpCAM | G8.8 | APC | eBioscience | 17-5791-80 |
| CD44 | IM7 | APC | BioLegend | 103011 |
| Ly-6G | 1A8 | APC | Biolegend | 127614 |
| CD3e | 145-2C11 | PE | Biolegend | 100307 |
| CD274 (PD-L1) | 10F.9G2 | APC | Biolegend | 124311 |
| CD103 | AF1990 | AF488 | R&D systems | FAB1990G |
| PVR | TX56 | APC | BioLegend | 131509 |
| CD86 | GL1 | APC | Biolegend | 105011 |
| CD279 (PD-1) | 29F.1A12 | APC | BioLegend | 135225 |
| CD335 (Nkp46) | 29A1.4 | BV421 | Biolegend | 137612 |
| CD8a | 53-6.7 | BUV395 | BD Biosciences | 563786 |
| CD206 (MMR) | C068C2 | APC | Biolegend | 141707 |
| CD4 | RM4-5 | FITC | Biolegend | 100509 |
| CD45R (B220) | RA3-6B2 | PE | Biolegend | 103207 |
| CD11c | N418 | APC | Biolegend | 117310 |

[1]target, marker that is recognised by the antibody; clone, antibody clone; fluorophore, fluorophore directly conjugated to the antibody; source; the company where the antibody was purchased, alongside the catalogue number.

All antibodies were tested in 1:40, 1:100 and 1:200 dilutions to assess sensitivity and confirm specificity.

### Supplementary Table 2. Validated Metal-conjugated IMC antibody panel

| metal[1] | target | clone | reactivity | source | catalogue number | dilution | purpose | cell type / phenotype | Used for clustering |
|---|---|---|---|---|---|---|---|---|---|
| 89Y | CD45 | 30-F11 | Mouse | Fluidigm | 3089005B | 1:50 | lineage | leukocytes | Y |
| 141Pr | alphaSMA | 1A4 | Mouse/human | Fluidigm | 3141017D | 1:100 | lineage/structure | fibroblasts | Y |
| 142Nd | MHC-II | M5/114.15.2 | Mouse | BioLegend | 107637 | 1:200 | lineage/phenotype | dendritic cells / macrophages | Y |
| 143Nd | vimentin | D21H3 | Mouse/human | Fluidigm | 3143027D | 1:100 | lineage | tumour cells | N |
| 144Nd | PECAM/ CD31 | 390 | Mouse | BioLegend | 102425 | 1:100 | lineage | endothelium | Y |
| 145Nd | | | | | | | | | |
| 146Nd | F4/80 | (CI:A3-1) | Mouse | BioRAD | MCA497GA | 1:100 | lineage | macrophages | Y |
| 147Sm | CD68 | FA-11 | Mouse | BioLegend | 137002 | 1:100 | lineage | macrophages | Y |
| 148Nd | | | | | | | | | |
| 149Sm | EpCAM | G8.8 | Mouse | BioLegend | 118223 | 1:200 | lineage | epithelium | Y |
| 150Nd | CD44 | IM7 | Mouse/human | Fluidigm | 3150018B | 1:100 | lineage | tumour cells / activation marker | Y |
| 151Eu | Ly-6G | 1A8 | Mouse | Fluidigm | 3151010B | 1:100 | lineage | neutrophils | Y |
| 152Sm | CD3e | 145-2C11 | Mouse | Fluidigm | 3152004B | 1:100 | lineage | T cells | Y |
| 153Eu | CD274 (PD-L1) | 10F.9G2 | Mouse | Fluidigm | 3153016B | 1:100 | phenotype | immune checkpoint | N |
| 154Sm | | | | | | | | | |
| 155Gd | CD103 | AF1990 | Mouse | R&D systems | AF1990 | 1:100 | lineage | dendritic cells | Y |
| 156Gd | | | | | | | | | |
| 158Gd | Foxp3 | FJK-16s | Mouse | Fluidigm | 3158003A | 1:100 | lineage | regulatory T cells | N |
| 159Tb | TCRgd | GL3 | Mouse | BD Biosciences | 553175 | 1:100 | lineage | gamma-delta T cells | N |
| 160Gd | PVR | TX56 | Mouse | BioLegend | 131502 | 1:100 | phenotype | immune checkpoint | Y |
| 161Dy | CD86 | GL1 | Mouse | BD Biosciences | 553689 | 1:100 | phenotype | costimulatory ligand | N |

| | | | | | | | | | |
|---|---|---|---|---|---|---|---|---|---|
| 162Dy | | | | | | | | | |
| 163Dy | | | | | | | | | |
| 164Dy | | | | | | | | | |
| 165Ho | | | | | | | | | |
| 166Er | CD279 (PD-1) | 29F.1A12 | Mouse | BioLegend | 135202 | 1:100 | phenotype | immune checkpoint | N |
| 167Er | CD335 (Nkp46) | 29A1.4 | Mouse | Fluidigm | 3167008B | 1:100 | lineage | NK cells | Y |
| 168Er | CD8a | 53-6.7 | Mouse | Fluidigm | 3168003B | 1:100 | lineage | CD8 T cells | Y |
| 169Tm | CD206 (MMR) | C068C2 | Mouse | Fluidigm | 3169021B | 1:50 | phenotype | M2/hypoxic macrophages | N |
| 170Er | pS6 | D68F8 | Mouse/human | NEB | 5364BF | 1:100 | phenotype | active mTOR signalling | N |
| 171Yb | CD4 | RM4-5 | Mouse/human | BioLegend | 100561 | 1:100 | lineage | CD4 T cells | Y |
| 172Yb | cleaved caspase3 | 5A1E | Mouse/human | Fluidigm | 3172027D | 1:100 | phenotype | apoptosis | N |
| 173Yb | Ki-67 | 16A8 | Mouse | BioLegend | 652402 | 1:100 | phenotype | proliferation | N |
| 174Yb | | | | | | | | | |
| 175Lu | | | | | | | | | |
| 176Yb | CD45R (B220) | RA3-6B2 | Mouse | Fluidigm | 3176002B | 1:100 | lineage | B cells | Y |
| 209Bi | CD11c | N418 | Mouse | Fluidigm | 3209005B | 1:100 | lineage | dendritic cells / macrophages | Y |

[1]Metal, the isotope to which the antibody was conjugated, note: antibodies purchased from Fluidigm, were bought in pre-conjugated format, antibodies from other suppliers were bought in a purified format and conjugated in house; target, marker that is recognised by the antibody; clone, antibody clone; reactivity, indicating the species to which the antibody is reactive to; source; the company where the antibody was purchased, alongside the catalogue number; dilution, the dilution at which the antibody was included in the antibody mix; purpose, whether the marker is used to define cell lineages or characterises a phenotype of the cell; cell type/phenotype, type of cell this marker is typically expressed on or functional meaning of the marker; used for clustering, whether this marker was selected for Phenograph clustering.

**Supplementary Table 3. Key markers to annotate clusters, related to heatmap Figure 3.**

| Cluster[1] | Counts | Cell type assigned | Key markers in order of importance for annotation (threshold where relevant) |
|---|---|---|---|
| 01 | 13681 | Tumour | $PVR^+$, $CD44^+$, $CD45^-$ |
| 02 | 15613 | Tumour | $PVR^+$, $CD44^+$, $CD45^-$ |
| 03 | 3586 | DC1 | MHC-$II^+$, $CD103^+$, $CD11c^+$, $CD45^+$ |
| 04 | 3341 | unclassified | Low on key markers |
| 05A | 3212 | Macrophages | $CD68^+$, $F480^+$(>0.5), MHC-$II^+$, $CD45^+$ |
| 05B | 4447 | Dendritic cells | $CD11c^+$(>0.6), MHC-$II^+$, $F480^{lo}$, $CD45^+$ |
| 05C | 28717 | Tumour | $CD44^+$, $PVR^+$, F4/$80^-$ |
| 06A | 1439 | Regulatory T cells | $Foxp3^+$(>0.6), $CD4^+$, $CD3^+$, $CD45^+$ |
| 06B | 6176 | CD4 T cells | $CD4^+$, $CD3^+$, $CD45^+$ |
| 07 | 2975 | Endothelium | $PECAM^+$ |
| 08 | 10857 | Macrophages | $F480^+$, $CD68^+$, $CD45^+$ |
| 09 | 5717 | B cells | $B220^+$, $CD45^+$ |
| 10 | 12827 | Tumour | $CD44^+$, $PVR^+$, $CD45^-$ |
| 11 | 10187 | Macrophages | $CD68^+$, $CD11c^+$, $CD45^+$ |
| 12 | 8465 | unclassified | Low on key markers |
| 13 | 6414 | Endothelium | $PECAM^+$ |
| 14 | 21197 | Endothelium | $PECAM^+$ |
| 15A | 14 | NK cells | $NKp46^+$(>1.5) , $CD45^+$ |
| 15B | 6024 | Tumour | $CD44^+$, $PVR^+$, $pS6^+$, $CD45^-$ |
| 16 | 12546 | Neutrophils | $Ly6G^+$, $CD45^+$ |
| 17 | 11445 | Tumour | $PVR^+$, $CD44^+$, $CD45^-$ |
| 18 | 1497 | Epithelium | $EPCAM^+$ |
| 19 | 5113 | Endothelium | $PECAM^+$ |
| 20 | 4458 | Neutrophils | $Ly6G^+$, $CD45^+$ |
| 21A | 918 | Macrophages | $CD206^+$(>0.5), $CD68^+$ (>0.5), $CD45^+$ |
| 21B | 3968 | Tumour | $PVR^+$, $CD44^+$, $CD45^-$ |
| 22 | 13633 | Tumour | $PVR^+$, $CD44^+$, $CD45^-$ |
| 23 | 230 | Macrophages | $F480^+$, $CD68^+$, $CD45^+$ |
| 24 | 5984 | Fibroblasts | $aSMA^+$ |
| 25 | 3097 | CD8 T cells | $CD8^+$, $CD3^+$, $CD45^+$ |
| 26 | 43433 | Macrophages | $CD206^+$, $F480^+$, $CD45^+$ |
| 27 | 5688 | DC other | MHC-$II^+$, $CD45^+$, $F480^-$ , $CD68^{lo}$ |
| 28 | 332 | Fibroblasts | $aSMA^+$ |
| 29A | 1051 | Epithelium | $EPCAM^+$ |
| 29B | 2199 | Endothelium | $PECAM^+$, $Vimentin^+$ |
| 30 | 2356 | Neutrophils | $Ly6G^+$, $CD45^+$ |

[1]Cluster, overview of clusters as obtained by Phenograph clustering and expert gating (for the clusters indicated with “A”, “B”, etc.), alongside the counts of cells within each cluster across the whole dataset. Cell type assigned, annotation given based on assessment of expression of key markers, in case of manual gating the thresholds are indicated.

**Supplementary Table 4. Linear regression analysis to rank changes in expression between treatment groups.**

Coefficients:

| | estimate | Std_error | t_value | p-value |
|---|---|---|---|---|
| MI_MHCcII | -0.1410488 | 0.00082112 | -171.77635 | 0.00E+00 |
| MI_CD86 | -0.1391809 | 0.00072477 | -192.03581 | 0.00E+00 |
| MI_F480 | -0.1385978 | 0.00070827 | -195.68529 | 0.00E+00 |
| MI_Vimentin | -0.1370623 | 0.0007154 | -191.58718 | 0.00E+00 |
| MI_CD206 | -0.1295906 | 0.00069023 | -187.75021 | 0.00E+00 |
| MI_CD45 | -0.1102835 | 0.00071768 | -153.66646 | 0.00E+00 |
| MI_aSMA | -0.0995303 | 0.00069105 | -144.02783 | 0.00E+00 |
| MI_CD44 | -0.0829157 | 0.00069306 | -119.63737 | 0.00E+00 |
| MI_CD68 | -0.0794998 | 0.00073016 | -108.87985 | 0.00E+00 |
| MI_PDL1 | -0.0629188 | 0.00077725 | -80.950323 | 0.00E+00 |
| MI_CD4 | -0.0534641 | 0.00059195 | -90.319095 | 0.00E+00 |
| MI_PECAM | -0.0528102 | 0.00080597 | -65.524162 | 0.00E+00 |
| MI_CD3 | -0.0515374 | 0.00060443 | -85.266268 | 0.00E+00 |
| MI_CD103 | -0.0423942 | 0.0008549 | -49.589724 | 0.00E+00 |
| MI_TCRgd | -0.0375577 | 0.00057673 | -65.12238 | 0.00E+00 |
| MI_casp3 | -0.0301824 | 0.0005335 | -56.574518 | 0.00E+00 |
| MI_PVR | -0.0280079 | 0.00081031 | -34.564538 | 3.04E-261 |
| MI_Foxp3 | -0.0229001 | 0.00055759 | -41.069736 | 0.00E+00 |
| MI_B220 | -0.0176966 | 0.00057821 | -30.605671 | 2.23E-205 |
| MI_CD11c | -0.0110315 | 0.00074868 | -14.734589 | 4.03E-49 |
| MI_Ki67 | -0.0082113 | 0.00062348 | -13.170065 | 1.34E-39 |
| MI_NKp46 | -0.0065885 | 0.00058215 | -11.317513 | 1.09E-29 |
| MI_CD8 | -0.0048249 | 0.00046561 | -10.362477 | 3.71E-25 |
| MI_LY6G | 0.02556174 | 0.00070279 | 36.371789 | 5.57E-289 |
| MI_EPCAM | 0.02730909 | 0.00076007 | 35.9298603 | 4.55E-282 |
| MI_PD1 | 0.03513414 | 0.00228591 | 15.3698457 | 2.74E-53 |
| MI_pS6 | 0.04067444 | 0.00075768 | 53.6832349 | 0.00E+00 |

Treatment group was one-hot encoded with “Vehicle” as 0 and “MRTX1257” as 1, to be the numeric predictor variable, and mean expression of each marker per cell was used in an independent linear regression analysis. Accumulated data was sorted by size of the coefficient estimate; negative values indicate a decrease and positive values an increase in expression related to MRTX1257 treatment.

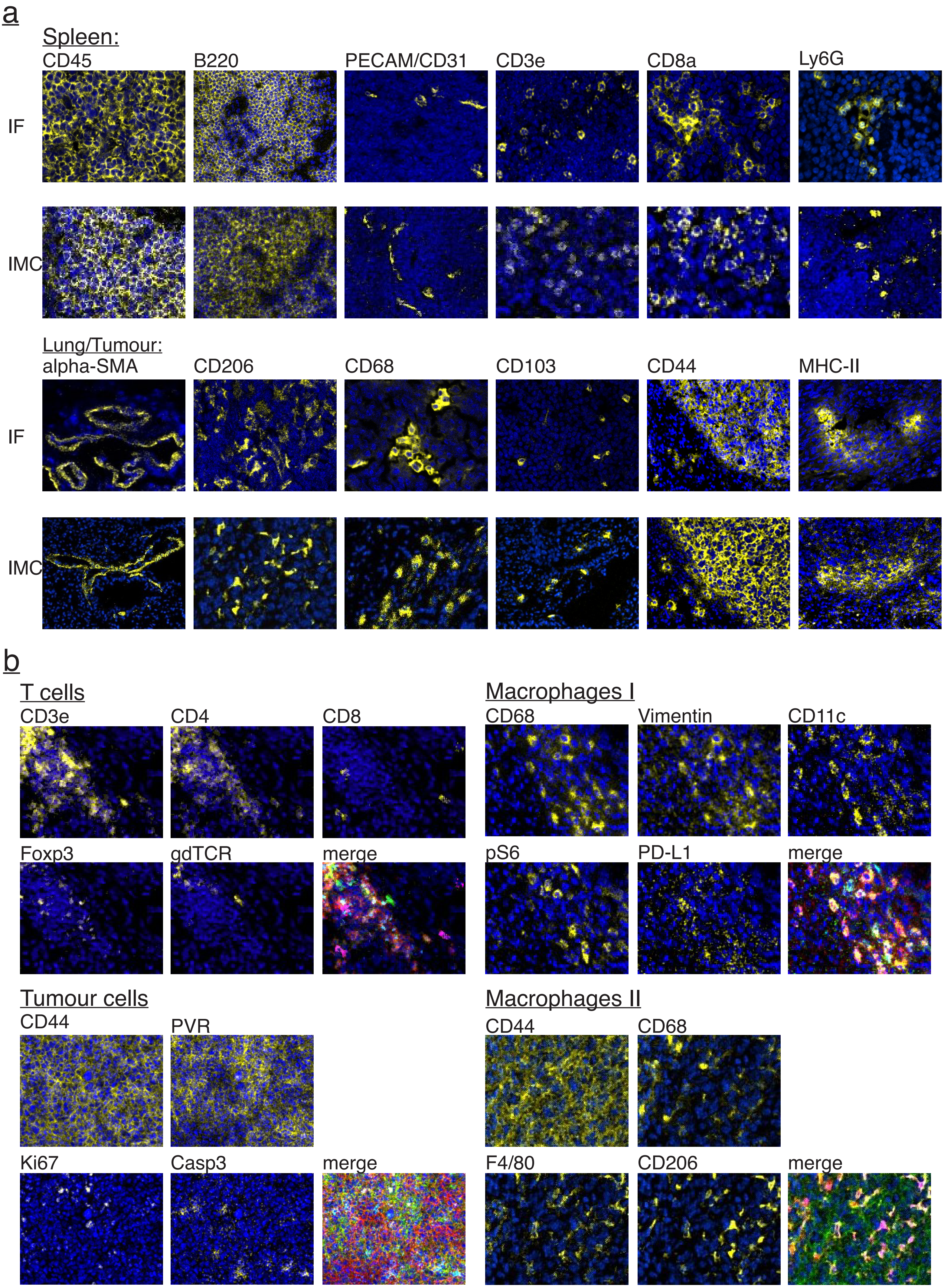


Supplementary Figure 1

**Supplementary Figure 1. Representative false colour images of markers used in the antibody panel.**

a. Mouse spleen and lung tumour tissue stained with fluorophore and metal conjugated antibodies. Images generated with either a Zeiss Upright 710 microscope with 20x objective lens or Hyperion Imaging Mass Cytometer. Most antibodies were tested in two independent stainings.

b. Lung tumour tissue of the 3LL ∆NRAS Lewis Lung Carcinoma model stained with metal-conjugated antibodies. Markers are grouped according to cell type, and each group contains a merged image showing co-expression of markers. Two independent stainings were performed to confirm findings.

Each marker is shown overlaid onto a nuclear marker (blue). Images were cropped, filtered with median filter, false coloured and individually scaled for visualisation in Fiji.

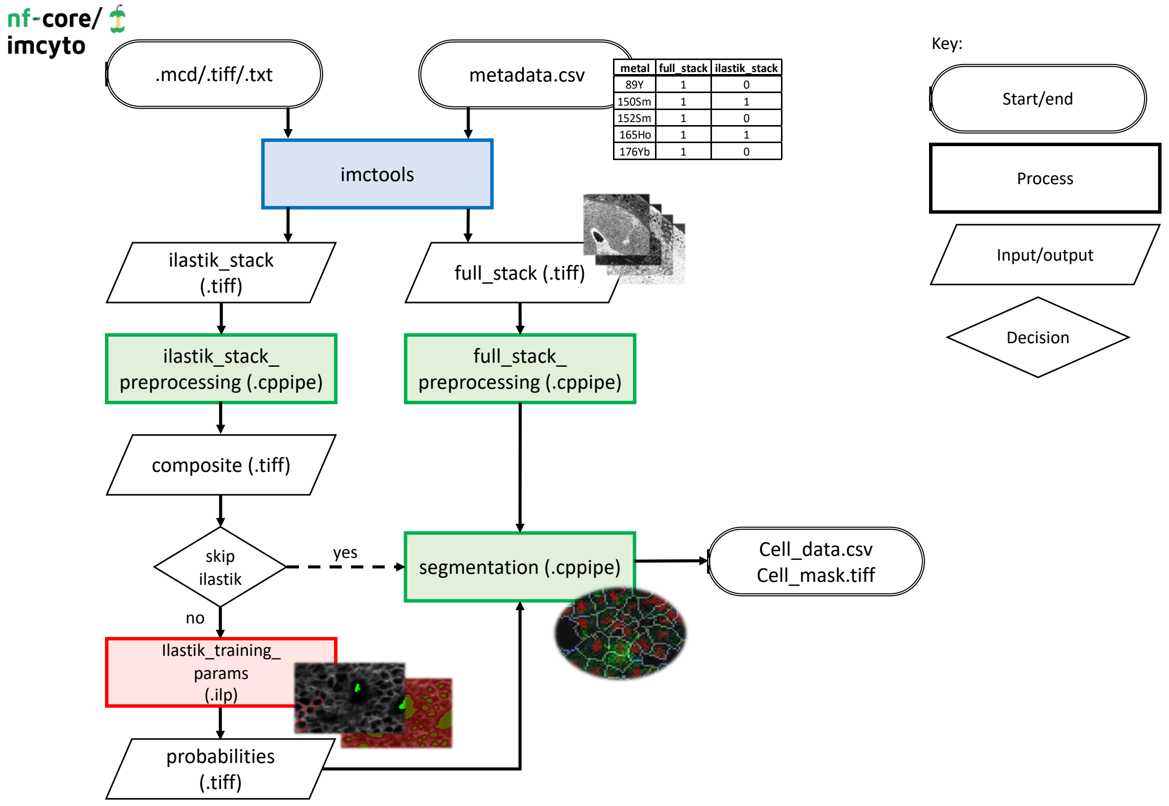


**Supplementary Figure 2. Schematic of the imcyto pipeline** (https://github.com/nf-core/imcyto or https://nf-co.re/imcyto). Raw data in the form of mcd/ome.tiff/txt files are taken as input alongside a metadata.csv file. The metadata.csv file contains a metal panel list with a Boolean tag (as shown) to determine which path each marker follows. Input data are converted into individual tiff images by imctools, matched against the metadata.csv and sorted into the desired path: either full_stack or ilastik_stack. Images to be used for Ilastik based image segmentation go down the ilastik_stack path to be pre-processed/filtered and undergo Ilastik pixel classification. Output probability maps are used to generate single cell masks in the segmentation step. The skip_ilastik parameter can be used to skip the Ilastik pixel classification altogether. Images in the full_stack path are minimally pre-processed and used in to extract single cell expression data from in the segmentation step. A single cell mask and csv file containing measured data are output at the end of the pipeline, to be used for further single cell analysis.

The key provided shows the input/outputs, process and decision steps in the flow diagram. Additionally, the process steps are colour coded by software used (blue-imctools, green-CellProfiler, red-Ilastik). The dashed arrow shows the workflow if skip_ilastik parameter is used.

Supplementary Figure 2

a b

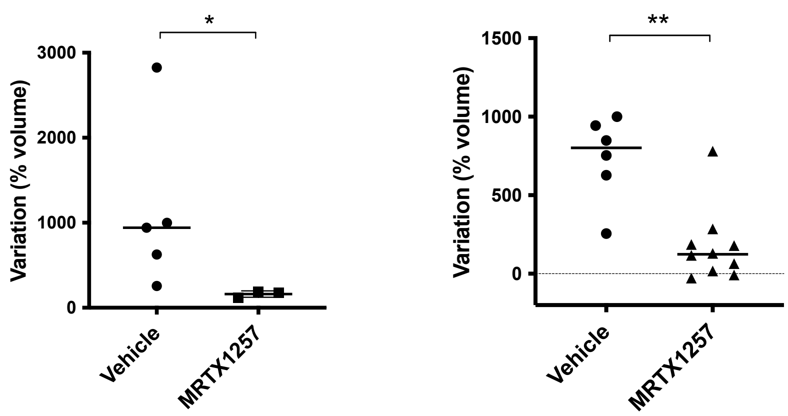


**Supplementary Figure 3. Lung tumour growth is inhibited following KRAS G12C inhibitor treatment.**

MicroCT analysis before and after 7 days of vehicle or MRTX1257 treatment in mice.

a. Tumour volume change per mouse, a sum of the individual tumours as depicted in b., from 5 vehicle treated mice and 3 MRTX1257 treated mice. * Significance: $P$ = 0.035 by a two-sided Mann-Whitney test.

b. Individual tumour growth, each dot or triangle represents the volume change of an individual tumour between the two time points; 6 tumours from 5 vehicle treated mice, and 10 tumours from 3 MRTX1257 treated mice. ** Significance: $P$ = 0.003 by a two-sided Mann-Whitney test.

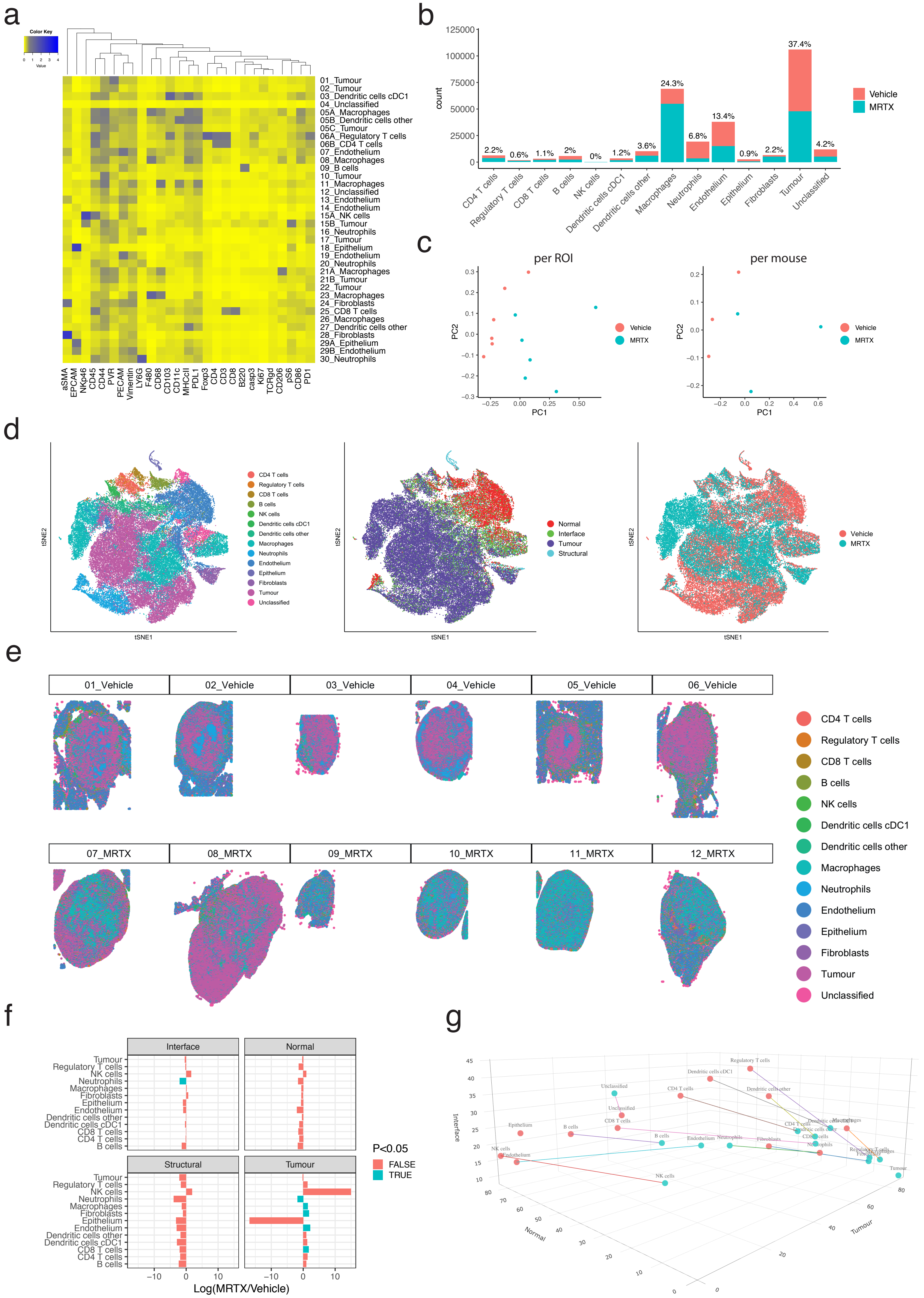


Supplementary Figure 4

**Supplementary Figure 4. Separation of cell types and ROIs by treatment**

a. Heatmap of all 36 cell types identified by Phenograph clustering and additional expert gating versus all markers in the panel. No scaling was applied to the heatmap.

b. Counts of cell types, coloured by treatment. Percentage on top of bars indicate the proportion of that cell type across the tissues.

c. Principal component analysis using the average of each marker per ROI (left) or per mouse (right) as input variables, coloured by treatment as indicated in the legend.

d. t-distributed stochastic neighbour embedding (tSNE). The tSNE plots were subsampled to 50.000 events for visualisation, but analysis was run on whole dataset. tSNE coloured by cell types (left), tissue domain (middle) and treatment (right).

e. *X*-*y* plots of the tissues coloured by cell type. Each cell in the dataset is plotted as a dot using its *x* and *y* coordinates in the ROIs and coloured by the cell type it was assigned to.

f. Relative distribution of cell types within the tissue domains compared between the two treatments, from an estimated marginal means calculation of a mixed effects model. Positive log(MRTX1257/Vehicle) values indicate an increased presence of that cell type in the respective domain in the MRTX1257 treatment group, negative values indicate increased presence in Vehicle treatment group. The larger the value, the bigger the differences between treatments.

g. Snapshot of 3D plot that shows the redistribution of cells across the tissue domains "Normal", "Tumour" or "Interface" as indicated on the *x*, *y* and *z* axes respectively. Per treatment (pink = vehicle, blue = MRTX1257), a dot marks the proportionate distribution (in percentage) of a cell type across these domains/axes. A coloured line connects the two treatment conditions to indicate the difference.

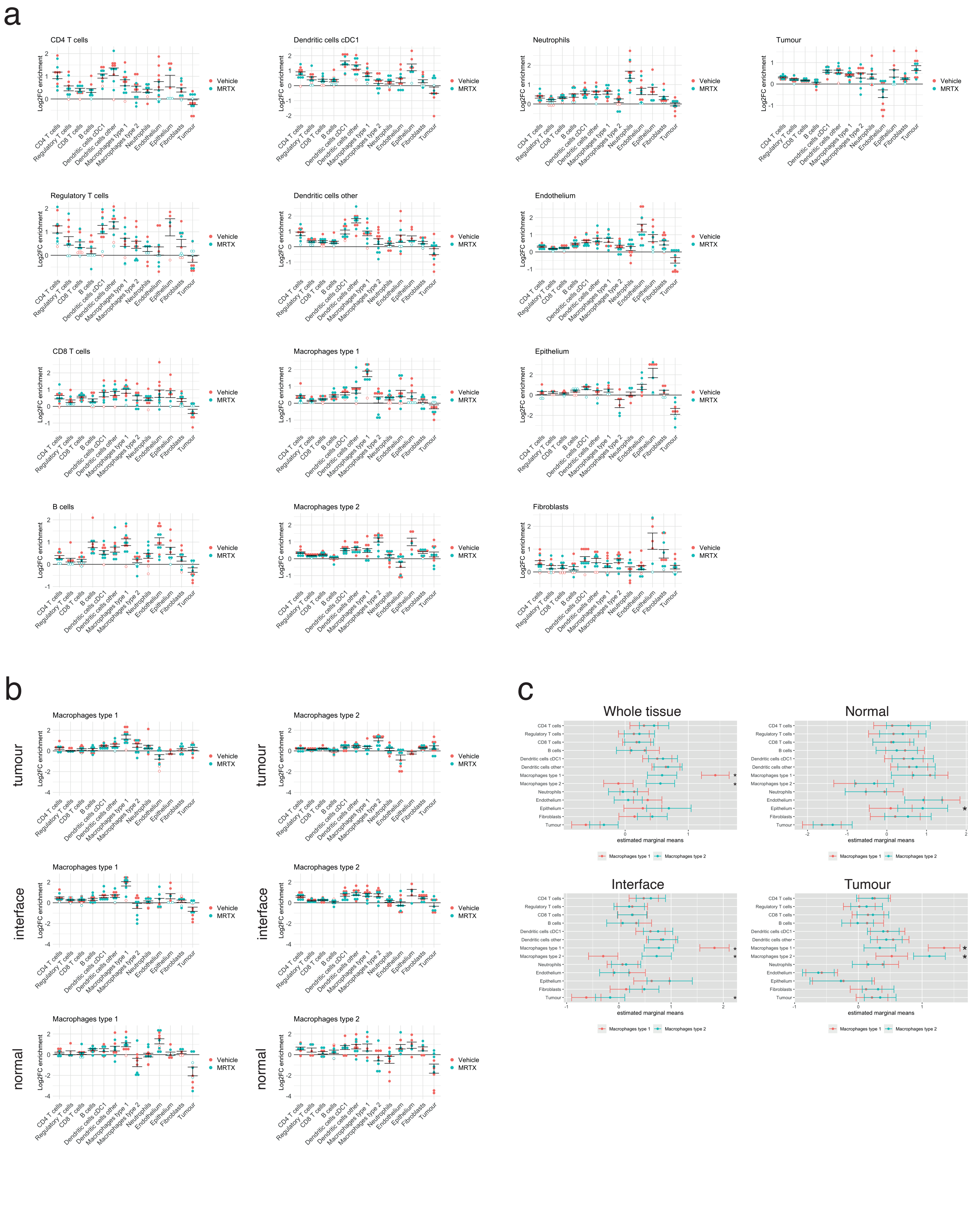


Supplementary Figure 5

**Supplementary Figure 5. Neighbourhood analysis**

a. “Log2FC enrichment” is the log2 fold change of the CT value for the frequency at which cell B is found in the neighbourhood of cell A, relative to the permutation test from the neighbouRhood[15] analysis (see Methods section). Six vehicle treated ROIs and 6 MRTX1257 treated ROIs were included in the analysis. Filled circles represent ROIs for which the enrichment was statistically significant ($p<0.01$), while open circles indicate non-significance. Where CT-real = 0 and CT-perm >0 this leads to an infinitely low log2FC value. Such Infs were omitted from the plots. Error bars indicate the standard error of means for the ROIs.

b. Log2 fold changes in neighbourhood enrichment as in a. for Type 1 and Type 2 macrophages in the different tumour domains as indicated. Filled circles represent images for which the enrichment was statistically significant compared to randomisation of all events in the image as calculated within the neighbouRhood package, while open circles indicate non-significance. Sample size: 6 vehicle and 6 MRTX1257 treated ROIs.

c. Depicting 95% confidence interval based on linear mixed-effects modelling to compare the neighbourhood enrichment plots from Supplementary Fig. 5b, incorporating the nested ROI-within-mouse variability, based on 6 ROIs from 3 vehicle treated mice and 6 ROIS from 3 MRTX1257 treated mice.

An asterisk indicates the cell types where the neighbourhood of Type 1 macrophages and Type 2 macrophages differ significantly for that domain, *P*-value <0.05 from estimating marginal means of the linear mixed-effects model.

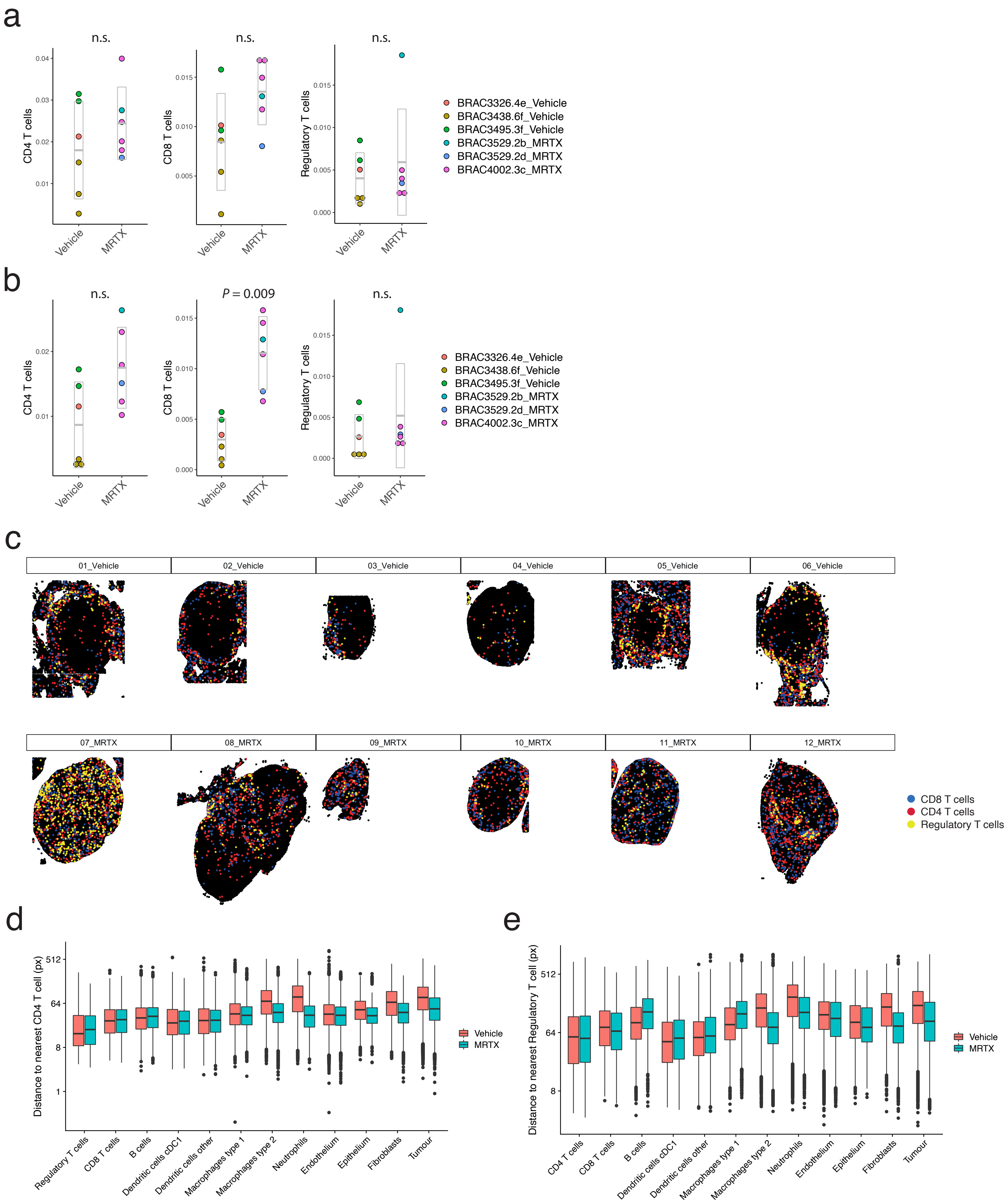


Supplementary Figure 6

**Supplementary Figure 6. T cells move into the tumour domain in response to MRTX1257**

a. Frequency of $CD4^+$, $CD8^+$ and regulatory T cells relative to total events within the ROIs. Depicting a total of 6 ROIs from 3 vehicle treated mice and 6 ROIS from 3 MRTX1257 treated mice, as coloured by mouse-ID. Differences between treatment groups are not significant according to linear mixed-effects model (estimated marginal means).

b. Frequency of $CD4^+$, $CD8^+$ and regulatory T cells relative to total events within the tumour domain of each ROI. Depicting a total of 6 ROIs from 3 vehicle treated mice and 6 ROIS from 3 MRTX1257 treated mice, as coloured by mouse-ID. *P*-values extracted from estimated marginal means of a linear-effects model.

c. *X*-*y* plots of tissues with T cells; $CD4^+$ T cells $CD8^+$ T cells and regulatory T cells in coloured dots as indicated, all other cells are shown in black.

d, e. Box-and-whiskers plots depicting the distance of cell types to the nearest $CD4^+$ T and regulatory T cell, for the whole single cell dataset (134186 cells from 6 Vehicle treated tumours and 148651 cells from 6 MRTX1257 treated tumours). Boxes minima and maxima represent $25^{th}$ and $75^{th}$ percentile, centre depicts the median, whiskers indicate 1.5*interquartile range, dots are individual outliers. Linear mixed-effects modelling confirmed that between treatments the cell types display a significantly different distribution of distances to nearest $CD4^+$ T cell ($P$ value = 0.020) and regulatory T cells ($P$ = 0.013, both for treatment:celltype interaction, ANOVA tested).

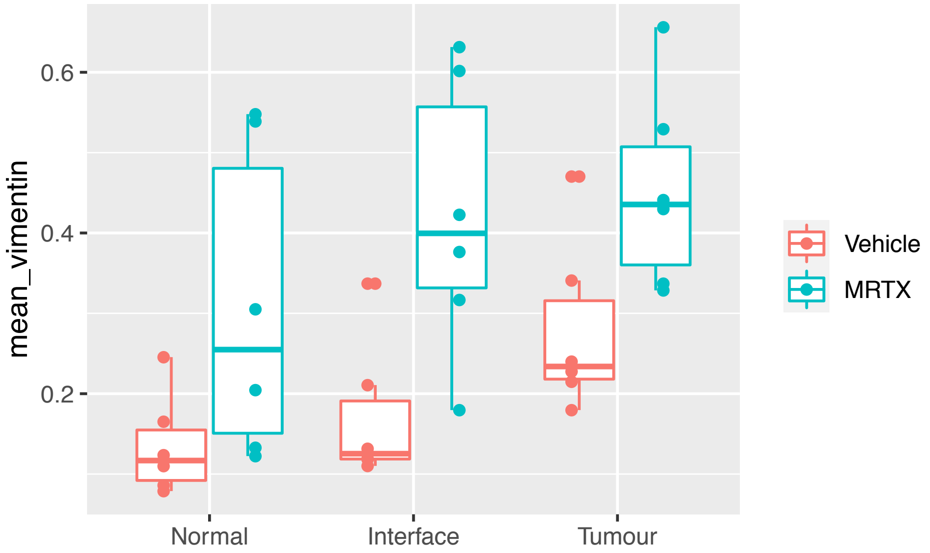


**Supplementary Figure 7. Vimentin expression in the tissue domains**

Box-and-whiskers plot depicting the median of vimentin expression on endothelium across the different tissue domains, boxes minima and maxima represent 25th and 75th percentile, centre depicts the median, whiskers indicate 1.5*interquartile range, sample size: 6 Vehicle treated tumours and 6 MRTX1257 treated tumours, a total of 37898 endothelium cells. Each point represents the mean intensity of vimentin expression on endothelium cells within the respective domain for one ROI.

**Supplementary Note 1. Interactive 3D visualisation of cell type distribution across domains**

The html file to load the Interactive 3D visualisation of cell type distribution across domains, related to the snapshot presented in Supplementary Fig. 4g, can be downloaded from https://hdl.handle.net/10779/crick.c.5270621. The proportions of the cell type within the normal, tumour and interface domains are used to position the cell types along the three axes of an interactive 3D plot, using the Plotly R Graphing Library. The data for the vehicle and MRTX1257 treated samples are connected with lines to emphasise the magnitude of changes in tissue distribution as a response to treatment.
